# A hypothetical new role for single-stranded DNA binding proteins in the immune system

**DOI:** 10.1101/320408

**Authors:** Nagarjun Vijay, Ajit Chande

## Abstract

The breadth of the host range of single-stranded DNA (ssDNA) viruses is roughly comparable to the host range of double-stranded DNA viruses (dsDNA). Yet, general ssDNA sensing receptors that activate the immune system have not been unequivocally identified while numerous dsDNA sensing receptors are known. Here, we hypothesize that some of the Single-Stranded DNA Binding (SSB) proteins may act as receptors that detect single-stranded DNA from pathogens and activate the innate immune system. As the first test of our hypothesis, we checked whether human genes that are known to bind to ssDNA are potentially interferon-regulated. Out of the 102 human genes that are known to have ssDNA binding ability 23 genes show a more than two-fold increase in gene expression upon interferon treatment. Single-stranded DNA viruses are pathogens of not only animals but also of plants and protozoans. We used this information to further prioritize our candidate list to ssDNA binding genes that are common between the model plant Arabidopsis thaliana and humans. Based on these strategies, we shortlist several promising candidate genes including the HMGB1 gene which could act as a ssDNA sensor that activates the immune system. Agreeably though we cannot establish a definitive role for these genes as ssDNA sensors of the immune system as yet, our preliminary analysis suggests the potential existence of ssDNA binding protein-like receptors (SLR’s) that are worth investigating further.

## Introduction

The innate immune system has been shown to consist of numerous receptors that are capable of recognizing pathogen associated molecules. These receptors are called pattern recognition receptors (PRRs) and the non-self-microbial molecules that these receptors recognize are called pathogen-associated molecular patterns (PAMPs). PRRs can detect a diverse set of ligands such as nucleic acids, proteins, carbohydrates, peptides, compound molecules, lipopolysaccharides, peptidoglycan, lipopeptides, phospholipids and glycoproteins (1, 2). After recognising the ligand molecule the receptors initiate downstream signalling to activate the immune response.

Within nucleic acid recognizing receptors, specific genes that detect ssRNA (single-stranded RNA), dsRNA (double-stranded RNA), CpG DNA, poly-nucleotide stretches, and DNA have been identified (3, 4). A considerable number of DNA sensing immune receptors have been identified in the past two decades (5). Receptors that specifically recognize ssDNA in a sequence independent manner have not been reported. However, certain ssDNA molecules that have an AT-rich stem-loop structure are thought to directly activate the immune response (6). Although it has been suggested that detection of ssDNA viruses might be possible through TLR9 (7) and IFI16 (8), it is unclear whether this is a general mechanism (9).

### The hypothesis

The Baltimore classification of viruses is based on the nucleic acid (RNA or DNA) used as the genetic material, presence of single or double strandedness, coding strand and mode of replication (10). Based on this classification system seven groups of viruses are recognised. Pathogen recognition receptors that are able to specifically identify dsDNA, dsRNA, ssRNA (both sense and antisense) have been experimentally characterised (3, 4). The gene TREX1 is able to degrade reverse transcribed DNA and the lack of the gene leads to build up of ssDNA generated by endogenous retro-elements (11). Infact, mutations in the TREX1 gene have been linked to initiation of autoimmune response (12, 13). The ability of TREX1 to degrade reverse transcribed DNA has been attributed to its exonuclease activity (14). Although TREX1 is the most highly up-regulated gene upon interferon treatment that also has ssDNA binding it cannot be considered to be an ssDNA sensor as it does not initiate a downstream response. The closest candidate for a true ssDNA sensor has been reported to be able to detect AT-rich ssDNA molecules (6). The well-known DNA sensing receptor IFI16 has also been proposed as a ssDNA sensor due to its ability to detect ssDNA stem loop structure (8). The existing literature on nucleic acid sensors has not been able to conclusively establish the existence of a true generic ssDNA sensor (15). It is possible that DNA sensors that specifically recognise the single stranded form of the DNA molecule don’t exist and the dsDNA sensors themselves are able to recognise ssDNA (16). However, this potential ability of dsDNA sensors to identify ssDNA molecules and initiating a immune response has also not been established (9).

Large scale profiling of meta-genomes has started revealing the true diversity of various classes of viruses by profiling previously inaccessible niches (17, 18). This has bolstered the idea that ssDNA viruses are probably more widespread among animal and plant species than previously estimated (19). New ssDNA viruses are also being discovered in host species such as archaea (20) and diatoms (21). The niches occupied by ssDNA viruses overlaps with that of other types of viruses enough for ssDNA viruses to have acquired genes from these other viruses through horizontal gene transfer to form a new family of ssDNA viruses (22). Since pathogen recognition receptors exist for every other class of viruses defined by the Baltimore classification, it seems unexpected that ssDNA sensing receptors have not been identified. If the ssDNA viruses were extremely restricted in their host range and/or ability to infect it could have explained this anomaly. However, the breadth of the host range of ssDNA viruses seems comparable to that of dsDNA viruses. This lack of general ssDNA sensors motivated us to hypothesize the existence of another class of PRRs that could act as receptors to specifically recognize ssDNA from pathogens such as ssDNA viruses. We call this hypothetical new class of receptors SLR’s (ssDNA binding protein-Like Receptors). Based on this hypothesis we make four predictions, (1) These receptors should possess ability to bind single stranded DNA, (2) Be regulated or responsive to the interferons or pro-inflammatory gene products, (3) These genes should be evolutionarily conserved at least across species that are permissive to the ssDNA viruses, and (4) Potentially be under episodic positive selection.

### Evaluation of the hypothesis – preliminary data analysis

To evaluate whether our hypothesis is a feasible scenario we searched public datasets and performed preliminary computational data analysis based on the four predictions from our hypothesis. We reasoned that an ssDNA sensor should probably have the ability to bind ssDNA. Genes with ssDNA binding ability have been assigned the Gene Ontology (GO) accession GO: 0003697. A total of 102 genes in the human genome come under this GO classification (see **Supplementary Table S1**). However, only 19 of these 102 genes have an oligonucleotide binding motif, or the OB fold. The OB fold (Interpro # IPR012340) is a five-stranded beta-barrel which is found in ss nucleic acid binding proteins. This OB fold structure is found in proteins with a wide range of functions such as DNA replication & repair genes (23).

The OB fold structure was absent from a majority of the genes with ssDNA binding activity we looked for other domains that are common among these genes. The P-loop NTPase fold (Interpro# IPR027417) domain was found in 20 of the genes. The P-loop NTPase fold catalyses NTP hydrolysis and utilises the energy released to induce conformational changes in other molecules. This catalytic activity makes genes with P-loop NTPase fold potential candidates for acting as receptors of ssDNA that possibly signal downstream molecules through conformational changes.

In addition to the ssDNA binding ability, pathogen recognition receptors should have the ability to increase their abundance upon a pathogenic attack to combat the greater number of pathogenic molecules. A vast majority of immune genes that are required to combat infection are regulated by a group of signalling proteins called interferons that are released in response to the presence of pathogens. These signalling proteins activate a group of transcription factor genes called signal transducer and activator of transcription (STAT) complexes and Interferon regulatory factors (IRF) to regulate the expression of various genes associated with the immune system (24). The complete set of genes that are directly induced by interferons or regulated indirectly is still being explored through genome-wide screens (25, 26). Publicly available gene expression datasets of interferon induced samples have been compiled and annotated to identify genes whose expression changes in response to interferon treatment. The Interferome database is an actively maintained comprehensive annotated collection of IFN-regulated genes (27, 28). We searched the Interferome database for the genes from our list of ssDNA binding proteins. However, presence of a gene in the Interferome database is just suggestive and not conclusive evidence for the gene being interferon regulated or not. Out of the 102 genes that are thought to have ssDNA binding ability, we found that 23 showed more than two-fold increase in gene expression upon interferon treatment. The type-1 interferon only regulation was seen for 4 genes, type-2 interferon only regulation was seen for 8 genes and 11 genes were found to be up-regulated independently by type-1 or type-2 interferon treatment (see **Supplementary Table S1** for details).

Pattern recognition receptors (PRRs) that detect DNA as well as those that detect RNA have been shown to be under positive selection (29–33), some have undergone recurrent whole gene or exon level losses and duplications (34–36). Prevalence of different types of pathogens has been a strong driver of evolutionary change in immune genes. The vast majority of genes that show strong signatures of adaptive evolution happen to be involved in the immune system in diverse species (37, 38) including humans (39). The hypersensitive response (HR) in plants is activated by resistance genes that code for receptor like proteins. Even these resistance genes found in host plants have been the targets of positive selection more often than other genes (40) and have been used to prioritise candidate resistance genes (41). Hence, genes that have experienced strong positive selection are likely to be involved in immune-related functions. This same line of reasoning has been used by previous studies to short-list immune related genes by looking at signatures of selection (42, 43). Only one of the 102 genes that have ssDNA binding ability is detected to be under positive selection in primates after using very conservative data analysis criteria (33). Surprisingly, we found that this gene contains a leucine-rich PPR motif (LR PPR Containing). The toll-like receptors that are involved in pathogen recognition for the innate immune system also have numerous successive leucine-rich repeat motifs but do not include the LRPPRC gene. Similarly, in plants, nucleotide binding site (NBS) -LRR gene family members are involved in activating the hypersensitive response (41, 44). The function of LRPPRC is only beginning to be understood and it has been implicated in suppressing autophagy. This gene has been implicated in suppressing autophagy and mitophagy by sequestering autophagy regulators Beclin 1 and Bcl-2 and the mitophagy regulator Parkin (45, 46). It has also been shown that the viral restriction factor Tetherin interacts with LRPPRC to activate autophagy (47).

Circular ssDNA viruses of the family Geminiviridae are considered to form a large fraction of plant pathogens (48). The flood of new data that is being generated from metagenomics screens has shown that circular ssDNA viruses have much greater diversity than previously estimated (19). However, plant genomes don’t have the homologous gene of LRPPRC. This could suggest the existence of other LRPPRC like genes as an alternative mechanism for ssDNA sensing in plants or that LRPPRC is not really involved in ssDNA sensing at all. Hence, as a fourth criteria we shortlisted genes with ssDNA binding ability in humans and the model plant *Arabidopsis thaliana*. We preferred to use a model plant species as it is likely to have more reliable annotation and would probably have been used for assigning GO terms in other species (49). Further restricting the gene set of 23 genes to those present in the plant *Arabidopsis thaliana* we are left with 2 genes (see **Supplementary Table S1**). We also choose few top candidates based on their similarity to known dsDNA sensors and ability to initiate a downstream immune response. In addition to these top candidates we also discuss few groups of genes that could also be potentially good candidates based on their conservation across animals and plants.

While experimental validation of the candidates would be a definitive test of the hypothesis proposed here, such experimentation is challenging following the existence of various other immune mechanisms for pathogen detection. Many of the candidates that we identify are no doubt involved in sensing Damage Associated Molecular Patterns (DAMP’s), however, the more important point that needs to be established is the ability of these sensors in being able to distinguish self from non-self-molecules. Whether the sensors are able to utilise the differences in base compositions of host vs pathogen DNA content in a spatio-temporal manner is an interesting idea worth exploring. Disentangling the role played by SLR’s if any is an arduous task. We simply provide a list of candidate genes which we prioritised based on the criteria described above. This gene list is not intended as definitive evidence for the existence of SLR’s. However, we hope that our hypothesis and preliminary reasoning based data analysis would motivate a more long-term search for the existence of SLRs.

### Empirical data – top candidate genes for single stranded DNA sensing

1. **HMGB1:** The HMGB1 gene has been the focus of numerous papers (50–54) and it has even been called the “nuclear weapon in the immune arsenal” (55). The role of HMGB1 in autoimmune diseases is also established (56). In addition to its role in the immune system, HMGB1 is also involved in regulation of gene expression (57). Studies in a animals and plants have established the HMGB family of genes as damage-associated molecular pattern (DAMP) molecules (58, 59). It should be also be noted however that the HMGB1gene has been proposed as a nucleic acid sensor that mediates an immune response (60, 61). However, a specific ability to detect various forms in which ssDNA could exist has not been explicitly discussed (62). HMGB1 has also been shown to have affinity for different DNA structures (52). Interestingly it has been shown to bind DNA mini-circles (63), looped structures (64) as well as hemi-catenated DNA (65) in addition to four-way junctions (66) and triplex DNA (67). HMGB1 is also one of the five genes in the human genome that has been assigned the GO term of bubble DNA binding (GO: 0000405). This broad range of DNA binding ability of HMGB1 and its established role in the immune response have not been linked with each other. Since the discovery of HMGB1’s role in DNA damage response has largely relegated it to the status of a DAMP (68). Single-stranded DNA viruses have many forms such as circular ssDNA found in many plant viruses as well as secondary-structures of linear ssDNA (69). It could be argued that the ability of HMGB1 to bind such a broad range of DNA structures makes it an ideal candidate to be an ssDNA sensor. In addition to being present in Interferome database, it has experimentally been shown that HMGB1 is actually interferon induced (70). Being a widely conserved protein it is present across animals, plants and protozoans. Given all these attributes HMGB1 would probably be the best candidate for being a SLR.
2. **<u>DDX11 & DHX9</u>:** Both these genes are part of the DEAD box protein family that is mostly involved in anti-viral innate immune response (71, 72). Contradictory to the role of other DEAD box genes, the DHX9 gene as well as DDX1 and DDX3 are known to be required for replication of numerous viruses (5). The retention of these genes that assists pathogens in the host genome is hard to explain. DDX3 is known to activate the immune system through MAVS (73). It has been suggested that these genes are potentially the target of the ongoing ‘arms race’ and have a role in innate immunity that might be exploited by pathogens (5). However, despite being part of the DEAD box protein family DDX11 has not been clearly linked to the innate immune response pathway.
3. **<u>RPA1, RPA2, RPA3 & RPA4:</u>** The Replication Protein A (RPA) is a protein complex involved in sensing DNA damage through association with ssDNA (74). It has already been proposed to activate a downstream immune response (75). All four RPA’s have ssDNA binding ability, but none of them are upregulated upon interferon treatment (see **Supplementary Table 1**).

### Other single stranded DNA sensor candidates

In addition to the top candidate genes discussed in the previous section, we believe the following set of genes could also be involved in innate immune sensing of single stranded DNA from pathogens. Based on their function and evolutionary relationships we have grouped these genes together.

1. <u>The Mini chromosome Maintenance protein complex (MCM)</u>: The MCM complex in eukaryotes has six genes, four of which are thought to be able to bind to ssDNA with only three of them having annotated OB fold domains. Only two of the six MCM’s with ssDNA binding ability showed a greater than two-fold increase in expression upon interferon treatment and also have annotated orthologs in A. *thaliana*. However, two other MCM genes show a more than two-fold decrease in gene expression. One of these two genes, MCM7 has been shown to play a role in the innate immune system through phagocytosis (76).
2. <u>Cold shock domain containing proteins:</u> The cold shock domain containing proteins YBX1 and YBX3 have the ability to bind ssDNA. While YBX3 gene showed a greater than two-fold increase in gene expression upon interferon treatment, YBX1 gene showed decrease in gene expression. However, a recent report has suggested that YBX1 might play the role of a receptor to detect bacterial cell wall components and activate the innate immune system (77).
3. <u>Nucleic Acid Binding Proteins (NABP):</u> The human genome has two genes in the NABP family. We see that NABP1 shows a greater than two-fold increase in expression upon interferon treatment while NABP2 does not. However, both genes contain the OB fold domain and are part of the Sensor of Single-Strand DNA Complex (SOSS Complex). The SOSS complex is a multiprotein complex involved in DNA repair and cell checkpoint activation and maintenance of genomic stability. NABP1 is also known to play an important role in regulating thymopoiesis and is expressed in immune cells (78). Since NABP’s can’t bind dsDNA they would provide specificity for ssDNA sensing.

### Consequences of the hypothesis

The evaluation of the hypothesis proposed here has far reaching implications. First, if the existence and identity of SLR’s are confirmed, this would greatly improve our understanding of the functioning of the innate immune system that integrates ssDNA sensing with immune activation. Signal initiators that act downstream from these receptors might reveal the existence of additional layers of regulation of the innate immune pathways. Many of the candidate genes that we identify as single stranded DNA sensors are part of the DNA damage response pathway. A similar pattern has been seen for dsDNA sensors that have been identified. Experimental validation of the hypothesised SLR’s could further reinforce the idea of co-option of the DNA damage response genes by the immune system (75).

One of our candidates for being an ssDNA sensor is the LRPPRC gene. A recently published study has identified 3 amino-acid residues (located at position 478, 889 and 1139) from the gene to be under positive selection among primates. All three of the positively selected residues are located outside the predicted LRR domains. The presence of positively selected residues outside the LRR domains in plant resistance genes has been interpreted as changes in pathogen recognition regions (41). Further investigation of the functional role of these positively selected residues would help clarify this. Another leucine rich repeat motif containing gene LRRFIP1 (Leucine-rich repeat flightless-interacting protein 1) has been shown to be a cytosolic nucleic acid sensor (79). LRRFIP1 activates the interferon response through its downstream partner beta-catenin. Similar to this dsDNA sensor our candidate gene LRPPRC could act as a receptor that activates downstream signalling through transcriptional regulation of MDR1 and MVP (80). The MVP gene codes for the Major Vault Protein. Multiple molecules of this protein come together to form larger sub-cellular structures called vaults. The function of these vaults has been a mystery despite the various apoptosis related functions that have been attributed to it (81). Recent work has established an important role for the Major Vault Protein in virus induced interferon response (82, 83). Hence, the interaction of LRRPC with MVP provides a potential new pathway for the activation of innate immunity.

The HMGB1 gene is well known for its role as a non-histone chromatin protein involved in gene regulation (84) and also as a mediator of immune response (85). Our preliminary analysis has suggested a yet another role for this HMGB1 gene as a single stranded DNA sensor. The downstream signalling cascade and initiation of immune response through the Toll-like receptor 4 as well as cGAS pathway have been characterised in detail (86–88). The fact that we find the HMGB1 gene from our screening of the databases could even be taken as evidence that our approach has probably been successful in identifying true SLR’s.

Among the various viral sensing mechanisms that have been discovered, Dicers acting through the RNA interference (RNAi) pathway are considered ancient enough to have existed in the LECA (Last eukaryotic common ancestor). The RIG-I-like receptors (RLRs) are a more recent innovation that is known to be present only in vertebrates. The Toll-like receptors (TLRs) potentially originated in the early Bilateria and are older than RLR’s (89). Interestingly, many plants have a greater diversity of genes in the Dicer pathway that are able respond to a diverse set of viral infections (90). Hence, the ancestral Dicer mediated anti-viral response of plants is much more powerful than that found in other eukaryotes.

If our hypothesis is wrong, it suggests the intriguing possibility that ssDNA sensing might be done by means other than a dedicated ssDNA sensing receptor. One possibility is that the immune system is directly activated by the recognition of the DNA damage response complex which is activated by ssDNA. This would require some way to distinguish self vs non-self ssDNA mediated DNA damage response. Another possible mode of ssDNA sensing could be through the formation of dsDNA by complementation of the ssDNA of the pathogen followed by recognition by dsDNA sensors. It has been shown that ssDNA viruses have used various mechanisms for insertion into their host genomes (91, 92). Transcription of such endogenous viral elements could lead to complementation of ssDNA and formation of DNA-RNA hybrids that would get detected by TLR9 followed by activation of the cGAS-STING pathway (93, 94). Single stranded DNA is always protected by the binding of ssDNA binding proteins. Hence, direct recognition of ssDNA might not be the best strategy for the detection of ssDNA from pathogens. Hence, it is possible that the single stranded DNA binding proteins or their complexes with ssDNA can be activators of the immune system. The Replication Protein A genes are promising candidates in this regard. In any case, this hypothesis provides interesting candidate single stranded DNA binding protein like receptor genes by novel thinking about pathogen recognition receptors (PRRs).

## Acknowledgement

NJ would like to acknowledge funding from IISER Bhopal under grant # INST/BIO/2017/019. AC is a recipient of Ramanujan Fellowship from the SERB, Department of Science and Technology, and an Innovative Young Biotechnologist Award from the Department of Biotechnology, (both Government of India). AC thanks IISER Bhopal for funding under grant # INST/BIO/2017/012.

